# The impact of turbidity on foraging and risk taking in the invasive Nile tilapia (*Oreochromis niloticus*) and a threatened native cichlid (*Oreochromis amphimelas*)

**DOI:** 10.1101/2020.07.23.217513

**Authors:** Jonathan D B Wing, Toby S Champneys, Christos C Ioannou

## Abstract

Anthropogenic activity can increase water turbidity, changing fish behaviour by reducing visibility. The spread of invasive species is also facilitated by human activity, further increasing the pressure on native species. In two experiments we measured the foraging efficiency, risk perception and inter-individual consistency of risk-taking (personality variation in boldness) of an invasive species, the Nile tilapia (*Oreochromis niloticus*), and a threatened tilapia, the Manyara tilapia (*Oreochromis amphimelas*), in clear and turbid water. In experiment one, *O. niloticus* was faster to initiate feeding, encountered more food items, and consumed more than *O. amphimelas*. The latency to start foraging by *O. niloticus* decreased in turbid water. Turbidity did not affect the latency to start foraging in *O. amphimelas* but the number of food items they encountered was highest at the intermediate turbidity. There was however no significant effect of turbidity in either species on the total food consumed. In contrast to this foraging context, in experiment two with a refuge and no food available, risk taking behaviour was similar in both species and they both responded with similarly reduced risk taking in turbid water. Evidence of personality variation was weak, being observed only in *O. amphimelas* when first leaving the shelter in turbid water. Overall, species differences were greater in the foraging context but turbidity was more important in the risk-taking context. *O. amphimelas* is more sensitive to turbidity during foraging, and *O. niloticus* is likely to have a competitive advantage in foraging situations, especially in degraded turbid habitats.

**Significance Statement:** Under human-induced environmental change, native species are often exposed to multiple stressors. Here we tested the responses of two cichlid fish to increasing turbidity. The Nile tilapia (*Oreochromis niloticus*), which is invasive throughout the tropics, and the Manyara tilapia (*Oreochromis amphimelas*), a threatened species, indigenous to Tanzania. We found that turbidity was beneficial to the foraging of *O. niloticus*, which in both clear and turbid water consumed and encountered more food than *O. amphimelas*. In contrast, without food present, both species displayed similar responses of increased risk perception in turbid water with little evidence of personality variation between individuals in either species. Our results suggest that invasive species tolerant of degraded habitats may outcompete less well adapted native species for food.

## Introduction

Turbidity is the cloudiness of water produced by suspended particles scattering light (Davies-Colley and Smith 2001). This alters the contrast between an object and its background, influencing the vision of aquatic organisms (Utne-Palm 2002). Many high productivity aquatic habitats are naturally turbid and are populated by species which are specifically adapted to these environments (Williams 1982; Fabricius et al. 2005; Van De Meutter et al. 2005; Wenger et al. 2011). However, numerous anthropogenic activities can rapidly increase turbidity above natural levels, resulting in up to a 10-fold increase in sediment loading (Hilton and Phillips 1982; Dearing and Jones 2003; Järvenpää and Lindström 2004; Erftemeijer et al. 2012). Sedimentation, the erosion and transport of particles, is a growing cause of habitat degradation in clear water systems and drives rapid changes to the sensory environment (Julien 1995; Mol and Ouboter 2004; Erftemeijer et al. 2012). Such changes influence the behaviour of many fish species that rely on vision, especially during foraging and predator-prey interactions (Guthrie and Muntz 1986; Abrahams and Kattenfeld 1997; Ferrari et al. 2010a; Wishingrad et al. 2015; Ehlman et al. 2020). As a result, the behavioural responses of species to increased turbidity can heavily influence the effect it has on the survival of individuals and populations (Caves et al. 2017).

The effect of turbidity on foraging efficiency varies across species and ecological contexts. In some cases, turbidity can increase the contrast of an object against its background, benefiting object detection and improving foraging (Hinshaw 1985). More generally however, turbid water has been shown to restrict the visual range of fish and reduce the efficiency of visual foragers (Utne-Palm 2002; Lunt and Smee 2015). This negatively affects prey detection, reducing the reaction distance within which predators can detect their prey (Barrett et al. 1992; Gregory 1993; Sweka and Hartman 2003; Quesenberry et al. 2007). Therefore, foraging success can be constrained even in areas of high prey density (Turesson and Brönmark 2007; Becker et al. 2016; Snow et al. 2018). This has been demonstrated in brown trout (*Salmo trutta*) and bluegill sunfish (*Lepomis macrochirus*) where foraging performance and reactive distance declined as turbidity increased (Vinyard and O’Brien, 1976; Stuart-Smith et al. 2004). Even after initial detection, turbidity may impact foraging, with some species displaying reduced attack success even in low levels of turbidity (8 Nephelometric Turbidity Units (NTU)) (Johansen and Jones 2013). A common behavioural response to reduced visibility is to increase activity, as it can increase encounter rates (Sweka and Hartman 2001; Granqvist and Mattila 2004; Harvey and White 2008). Despite this, such behaviours are potentially detrimental if improved encounter rates do not compensate for the increased cost in energy or the greater risk of encountering predators (Meager and Batty 2007).

Turbid environments also influence predator-prey interactions by changing the perception of risk in fish species (Gregory 1993; Chamberlain and Ioannou 2019; Ehlman et al. 2019). Chinook salmon (*Oncorhynchus tshawytscha*) reduce antipredator behaviour in turbid conditions, suggesting a decreased perception of risk (Gregory 1993). This can be maladaptive if the true risk of predation does not decrease (Abrahams and Kattenfeld 1997; Lehtiniemi, et al. 2005). Additionally, turbidity can impair the ability of some fish to learn about and recognise predators, further slowing the reaction time of prey (Ferrari et al. 2010b; Chivers et al. 2013). However, in other cases turbidity reduces activity and feeding while increasing shelter use, indicating an increase in risk perception (Leahy et al. 2011; Ajemian et al. 2015).

Changes to foraging efficiency and risk perception due to turbidity thus impact the fitness of fish species (Tuomainen and Candolin 2011). However, individuals within a population may react differently to similar stimuli and this inter-individual consistency of behaviour is referred to as animal temperament or personality variation (Réale et al. 2007). A behavioural trait that is often consistently different between individuals is boldness, defined by the likelihood of an individual to take risks in return of greater rewards (Réale et al. 2007). The repeatability of behaviours associated with boldness can be induced by the combined effect of turbidity and the presence of predators (Ehlman et al. 2019). As personality variation indicates a lack of behavioural plasticity, it suggests that individuals may be constrained in their response to environmental change (Tuomainen and Candolin 2011).

In addition to turbidity, the introduction of non-native species poses a great threat to aquatic ecosystems, reducing biodiversity by displacing native species and contributing to habitat degradation (Mainka and Howard 2010). The casual mechanisms underlying the success and negative effects of many invasive species are, however, poorly understood (MacDougall and Turkington 2005; Tuomainen and Candolin 2011; Gallardo et al. 2016). Improving understanding of these mechanisms is an important goal of invasive species research, as it is crucial in making informed decisions regarding environmental management, and when allocating limited resources to the prevention of the most harmful species (Vander Zanden and Olden 2008). Successful invasive species often exhibit tolerance to a wide range of environmental conditions and adapt quickly to novel environments. This characteristic is so prevalent that the presence of invasive species has been suggested as an indicator of habitat degradation (Kennard et al. 2005; García-Berthou 2007). Improved tolerance of degraded environments can provide a competitive advantage for non-native species, allowing effective colonisation as intolerant native species decline (MacDougall and Turkington 2005; Linde et al. 2008a). For example, the combination of turbidity and the presence of the invasive yellowfin shiner (*Notropis lutipinnis*) reduces the feeding efficiency of the royside dace (*Clinostomus funduloides*) from the combined effects of restricted reaction distance and interspecific competition (Hazelton and Grossman 2009a, b). Resident species are often exposed to multiple stressor effects from anthropogenic activity (Ormerod et al. 2010; Orr et al. 2020), which can have a greater combined effect than each stressor in isolation (i.e. a synergistic effect).

The Nile tilapia (*Oreochromis niloticus*) is a widespread invasive cichlid species commonly used in aquaculture. Subsequently, non-native populations of *O. niloticus* have been established across the tropics (Canonico et al. 2005; Mohamed and M Al-Wan 2020). Like many invasive species, *O. niloticus* shows a high tolerance to a wide range of environmental conditions. For example, juvenile *O. niloticus* abundance has been shown to increase degraded habitats, while the abundance of native species declines (Linde et al. 2008b). Additionally, *O. niloticus* can actively increase turbidity through benthic foraging, by resuspending sediment and increasing nutrient levels leading to algal growth (Zhang et al. 2017). In this experiment, we investigated the behavioural responses of *O. niloticus* and the Manyara tilapia *Oreochromis amphimelas*, in clear compared to turbid water. *O. amphimelas* is a threatened tilapia species native to Tanzania where it lives in sympatry with the invasive *O. niloticus* across five lakes (Shechonge et al. 2019). In both species, we measured how foraging efficiency changes across low, medium, and high turbidity (0, 15, and 30 NTU), and whether risk taking and consistent inter-individual differences in risk taking (i.e. personality variation in boldness) were different in clear and turbid treatments (0 and 15 NTU). By investigating the relationship between turbidity and the behaviour of native and invasive fish species, the experiment aimed to provide insight into the behavioural responses of two ecologically relevant species with potential consequences for native populations that are exposed to the stressors of both increasing turbidity and invasive species.

## Materials and Methods

### Subjects

Sixty-eight *O. amphimelas* (65.3 ±□7.5 mm mean ± S.D body length) were bred from first generation wild caught stock by Bangor University and transported to Bristol in December 2018. Sixty-eight *O. niloticus* (79.6 ±□6.9 mm mean ± S.D body length) were purchased from Fish Farm UK, London, in December 2018. Thirty-six *O. amphimelas* and 36 *O. niloticus* were used in experiment one and 29 *O. amphimelas* and 32 *O. niloticus* were used in experiment two. Individuals were not reused between the two experiments and fish were housed in separate tanks according to the experiment they participated in. Three *O. amphimelas* were removed during the second experiment due to ill health during the trial period. The two species were housed separately in clear water (0 NTU) within a recirculating aquarium system in 180 litre glass tanks. Plastic plants and pipes were placed in the housing tanks to provide enrichment (Brydges and Braithwaite 2009). Identical shelters used in the experiments were also placed in the housing tanks to reduce novelty in the experimental trials (Fig. 1c). Fish were fed daily with a diet consisting of ZM large Premium Granular feed (Techniplast, London, UK), TetraMin flake (Tertra, Melle Germany), frozen bloodworm (CC Moore & Co, Templecoombe, UK) and Gamma ™ Krill Pacifca, chopped prawn, Mysis shrimp, Brineshrimp, and vegetable Diet (Tropical Marine Centre, Chorley wood, UK). Water temperature in the housing tanks was sustained at 26°C with a constant diurnal light cycle of 12:12 hours (light: dark) mimicking tropical conditions. All fish remained in the laboratory for future use following the experiments.

**Fig. 1.**
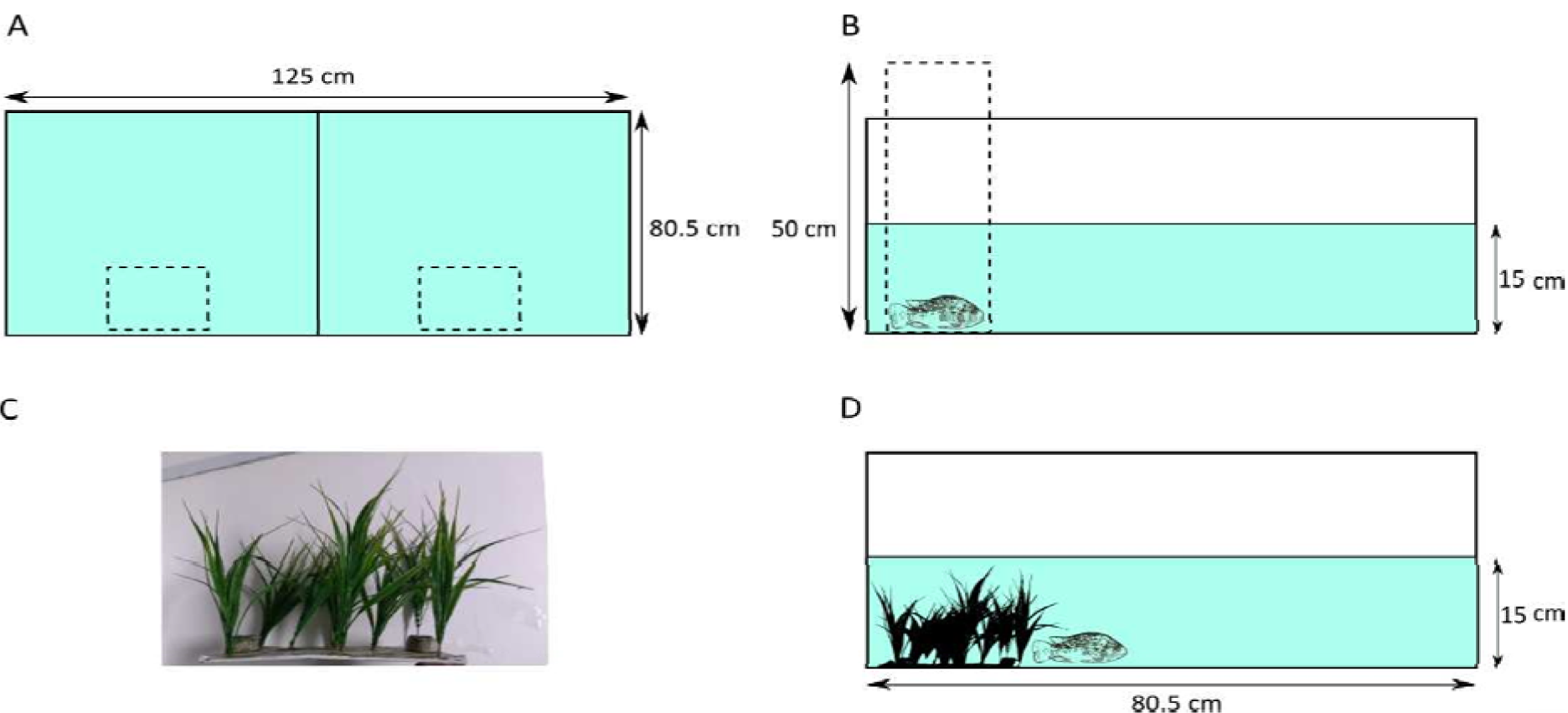
The experimental arena and shelter used in both experiments (not to scale). (A) Top-down view of the experimental arena and the position of the acclimation boxes used in experiment one (dashed). (B) Side view of the trial arena, with the removable acclimation box (dashed). (C) The plant shelter used in experiment two, made from plastic aquarium plants fixed to a plastic base and weighted with ornamental rocks. (D) Side view of the trial arena showing the position of the plant shelter used in experiment two where the fish were placed at the beginning of the trial.

### Experimental setup

The trials were conducted in a white acrylic tank with an opaque divider creating two arenas (each 80.5 × 64 × 40cm (width x length x height) containing 77.2 litres of water at a depth of 15cm) so that two trials could be conducted simultaneously (Fig. 1a). A shelter created from plastic ornamental plants attached to a white corrugated PVC base was placed in each side of the tank to act as a shelter in experiment two (Fig. 1c). Turbidity was recorded with a calibrated Thermo Scientific™ Orion™ AQUAfast AQ3010 turbidity meter□using□Nephelometric□Turbidity□Units (NTU). The tank was lit from above with a fluorescent strip light and recorded from above with a□GoPro Hero 5 Black (GoPro, Inc.) at a resolution of 1920 × 1080 pixels. The tank was filled with water from the recirculation system used to house the fish. A white curtain was drawn across the tank area to prevent disturbance.

### Procedure

#### Experiment one: The effect of turbidity on foraging efficiency

Each day of testing, one of three turbidity levels (0, 15 and 30 NTU) was randomly selected and 12 individuals (six of each species) were tested. Kaolin clay was used to create treatments of 15 NTU (0.04 mg/l) and 30 NTU (0.09 mg/l). The clay powder was spread evenly across the tank and mixed until the desired level of turbidity was attained following previous studies (Leahy et al. 2011; Quesenberry et al. 2007). The turbidity range used was similar to levels in other studies, staying below the upper tolerance levels of fish and not interfering with the transmission of chemical cues or altering the pH or hardness of the water (Horppila et al. 2004; Leahy et al. 2011). Anthropogenic activities such as mining can result in increasing levels of clay entering aquatic systems, making clay an ecologically relevant way of replicating natural levels of turbidity (Eriksson et al. 2004; Kemp et al. 2011). Resuspension by stirring was conducted between trials to ensure that the turbidity remained at desired levels across all trials.

The day before each trial, 6 individuals of each species were caught haphazardly with a hand net from their housing tanks and acclimatised to the next day’s turbidity (0, 15 or 30 NTU) in the trial tank for 16 hours (overnight) prior to testing. The two species were acclimated separately in the two halves of the tank, separated by the opaque divider. Fish were not fed the day before the experiment to encourage feeding and standardise hunger and motivation. Before the first trial on each day, all fish were removed and kept in covered plastic holding tanks (45 × 32 × 25cm, water depth 15cm) separated by species and with a turbidity matching that day’s treatment. One *O. niloticus* and one *O. amphimelas* were netted, their total body length measured (mouth to the end of caudal fin) with digital callipers, and were placed in an acclimation box in the trial area for 30 minutes before trials began (Fig. 1b). After this period, 10 pellets were added to each side of the experimental arena, spread evenly across the width opposite to the acclimation boxes. Floating pellets were used as food items (Hikari Cichlid Gold mini-pellet, 3.2-3.7mm) as they remained at the water surface and feeding attempts could be easily observed even in turbid conditions. Recording began and the acclimation boxes were slowly lifted and removed, releasing the fish. Trials lasted 15 minutes. Each fish was tested once and then placed into housing tanks reserved for fish who had completed the trials to avoid repeated testing of the same individual. The arenas and holding tanks were emptied after each day of trials and replaced with fresh system water which was dosed with the desired turbidity for the next day. The next group of fish was then acclimated for the next day’s trial. Trials took place between 30.04.2019 and 14.05.2019.

Video recordings were analysed using BORIS version 7.7.3 by a single reviewer to ensure consistency (Friard and Gamba 2016). The variables measured were the time to first feed, the number of feeding attempts (the number of the 10 pellets that the fish attempted to feed on) and the number of pellets consumed. A feed attempt was defined as an individual’s mouth making contact with a pellet at least once, so that subsequent attempts to feed on that pellet were not further counted. The number of feed attempts thus measures the number of different food items that were encountered during the trial. It was not possible to record data blind as individuals had to be identified by species for each experiment.

#### Experiment two: The effect of turbidity on risk taking and personality variation

Consistent inter-individual differences in responses to turbidity were tested by exposing each fish to each treatment (0 and 15 NTU) twice across a 4-day period. Eight individuals of each species were tested per week and every individual took part in one trial per day for four consecutive days. Turbid water was created following the methods outlined in experiment one. The day before each 4-day trial period, 8 individuals of each species were caught haphazardly with a hand net. Fish were split into 4 groups (2 groups of *O. niloticus* and 2 groups of *O. amphimelas*) and the total body length of each fish was measured (mouth to the end of caudal fin) using digital callipers. Each group was housed in a plastic holding tank (45 × 32 × 25cm, water depth 15cm), which were continuously oxygenated with air stones, for the next 4 days. These holding tanks were filled with system water which was changed each day with a turbidity level to match the next day’s trials (0 or 15 NTU), using the methods outlined in the previous experiment, allowing the fish to be acclimated for 16 hours (overnight) before the experiments began in the required treatment. The fish in each holding tank were different lengths to ensure clear identification of each individual was possible across the 4-day period. Trials took place between 18.06.2019 and the 26.07.2019.

The order of testing for groups, and individuals within groups, was allocated randomly per day. The water was stirred before each trial to maintain turbidity levels. Before each trial the turbidity was measured to make sure it was at the desired level (0 or 15 NTU) or resuspended by stirring if needed. Video recording began and individuals were placed within the shelters (Fig. 1c, d). Trials lasted for 30 minutes. Once tested, fish were returned to their holding tank. To standardise hunger levels between trials, fish were only fed after all individuals had taken part in the day’s experiments. At the end of the 4-day trial period, test fish were removed from the holding tanks and kept separately to untested fish to avoid individuals being used more than once. BORIS version 7.7.3 software (Friard and Gamba 2016) was used to analyse the videos by a single reviewer to maintain consistency. The variables measured were the time to first leave the shelter, the time taken to first cross the midline of the tank after first leaving the shelter, and the total time spent out of the shelter. It was not possible to record data blind because our study required individuals to be identified across repeat experiments.

### Statistical analysis - Experiment one

R version 3.5.3 was used for the analysis of both experiments (R Core Team 2019). The time to first feed, a censored time-to-event response variable, was analysed using a Cox proportional hazard test. The covariates included for this analysis were turbidity, species and body length. The principal assumption of this test is that the hazards are proportional, meaning that the effect of covariates do not change over time. The proportional hazards assumption was tested using Schoenfeld residuals (from R function “cox.zph” in the “survival” package) (Therneau 2015). To test the assumption of non-linearity, the martingale residuals were tested (from R function “ggcoxfunctional”, in the “survminer” package) (Kassambara et al, 2019). The deviance residuals were used to examine influential observations (from R function “ggcoxdiagnostics”, in the “survminer” package) (Kassambara et al, 2019). We found no evidence that these assumptions were violated. Additional models were also run on the data subset by species to determine the species-specific effects of turbidity on the response variables.

The number of feed attempts was analysed with negative binomial GLMs (generalised linear models). The continuous variables were scaled on all models (R function “scale”) and the dispersion parameter was calculated to ensure it was between 0.5 and 2 indicating no overdispersion (Duffield and Ioannou 2017). The number of feed attempts model initially included an interaction term between species and turbidity and their main effects, along with a continuous variable (body length), and a categorical variable (the side of the arena that the trial took place in). After observing a nonlinear relationship between turbidity and the number of feed attempts, polynomial regression was applied to provide a nonlinear fit to the model for these models (R function “poly”) (Becker et al. 1989; James et al. 2000; Chambers and Hastie, 2017). Likelihood Ratio Tests were used to remove any nonsignificant interactions or explanatory variables by deleting terms based on likelihood ratio tests (using R function “drop1” in the “lme4” package) (Bates et al. 2015).

The number of pellets consumed was analysed with negative binomial GLMs. The number of pellets consumed model initially included an interaction term between species and turbidity and their main effects, along with a continuous variable (body length), and a categorical variable (the side of the arena that the trial took place in). Additionally, to investigate the effect of turbidity within species, negative binomial models were created for each species separately. These included the main effect of turbidity along with continuous (body length) and categorical (the side of the arena that the trial took place in). predictor variables. Continuous variables were scaled and assumptions were tested using the same methods as outlined for the number of feed attempts. Once again, the “drop 1” function was used to remove nonsignificant interactions and explanatory variables.

### Statistical analysis - Experiment two

The time taken to first leave the shelter and the time taken to first cross the midline of the arena, censored time-to-event response variables, were analysed using Cox proportional hazards models. The covariates included for this analysis were turbidity, species and body length. The assumptions of the tests were tested following methods detailed in experiment one and the assumptions were met in all cases. Additional models were run on the data subset by species to determine the species-specific effects of turbidity on the response variables.

The total time spent outside the shelter was analysed with a generalised linear mixed model (GLMM) with a Poisson distribution. The model included an interaction between species and turbidity and their main effects, a continuous variable (body length) and a categorical variable (the side of the arena that the trial took place in) and a random effect (fish ID). Additional models were run on the data subset by species, and these models included turbidity along with categorical (the side of the arena that the trial took place in) and continuous (body length) variables and a random effect (Fish ID). To reduce overdispersion, continuous variables were scaled and an observation-level random effect term was included in the model (Harrison 2014). Likelihood Ratio Tests were then used to remove non-significant interactions by deleting terms based on chi squared tests using the “drop 1” function.

To assess consistency in inter-individual variation (i.e. personality variation) for each of the three response variables, in each species and in each of the turbidity treatment separately, Spearman rank non-parametric correlations were used. Shapiro-Wilk normality tests showed the data was not normally distributed (*P* < 0.05) and the censored data for the two latency response variables can lead to spurious estimates of formal repeatability scores (Stamps et al. 2012; Ioannou and Dall 2016).

## Results

### Experiment one

The likelihood of the first feeding attempt was significantly lower in *O. amphimelas* than *O. niloticus* across all treatments of turbidity (Cox Proportional hazard model (CPH): hazard ratio (HZ) = 0.21, N = 72, *P* < 0.001; Fig. 2).

**Fig. 2.**
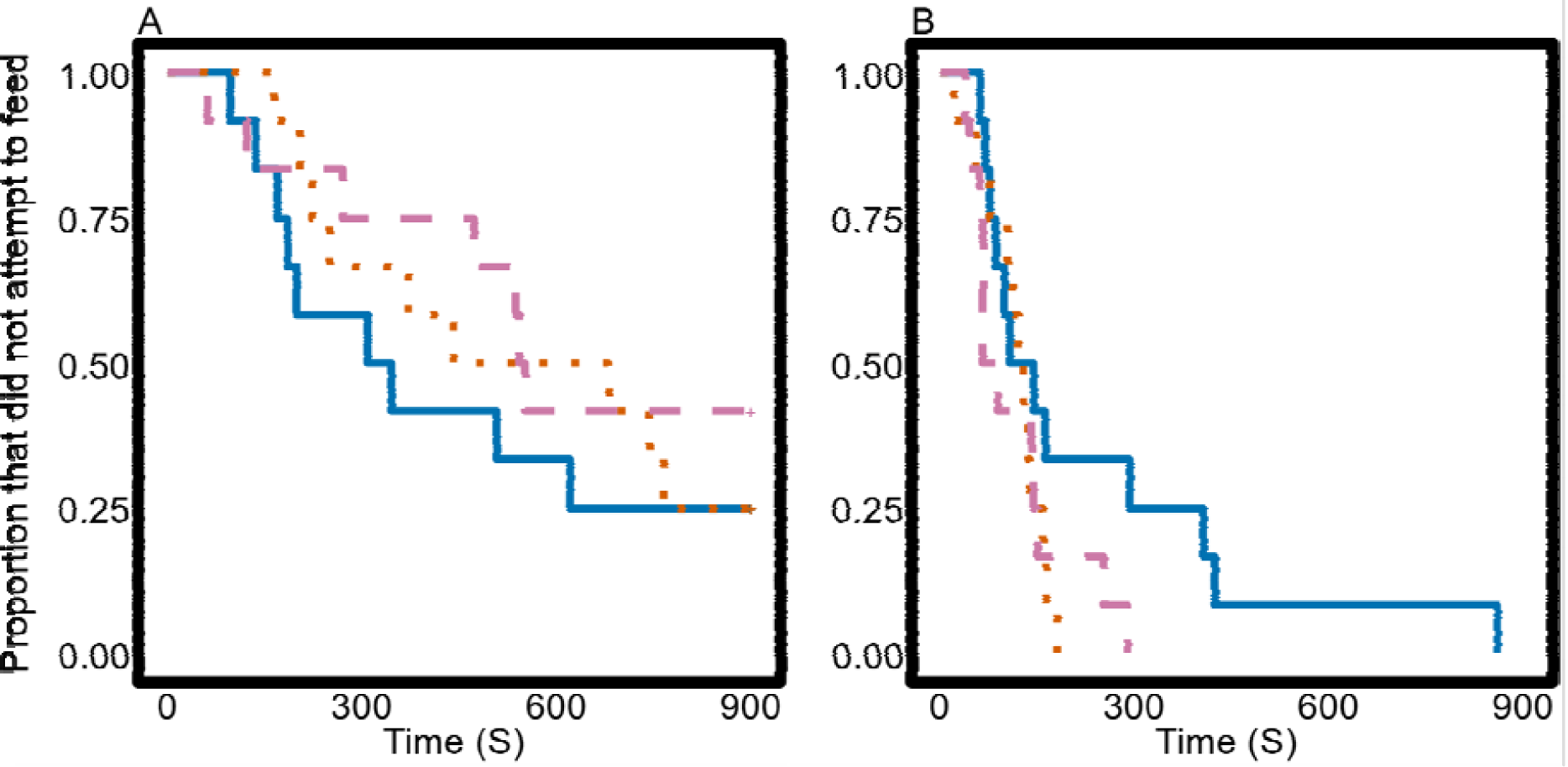
Kaplan-Meier estimates of the proportion of fish that did not feed across the trial period (A) *O. amphimelas* and (B) *O. niloticus* in 0 (solid), 15 (dotted) and 30 (dashed) NTU treatments.Turbidity did not affect the likelihood of the first feeding attempt in *O. amphimelas* (0 NTU versus 15 NTU CPH: HZ = 0.8, N = 36, *P* = 0.64; 0 NTU versus 30 NTU, CPH: HZ = 0.58, N = 36, *P* = 0.29; 15 NTU versus 30 NTU CPH: HZ = 0.72, N = 36, *P* = 0.53; Fig. 2a). However, *O. niloticus* was more likely to first feed at 15 and 30 NTU than at 0 NTU (0 NTU versus 15 NTU CPH: HZ = 2.5, N = 36, *P* = 0.04; 0 NTU versus 30 NTU CPH: HZ = 2.29, N = 36, *P* = 0.005; Fig. 2b). The likelihood of a first feed attempt did not differ between 15 and 30 NTU (CPH: HZ = 0.91, N = 36, *P* = 0.83).

When the model was restricted to fitting a linear relationship between turbidity and the number of the food items that the fish attempted to feed on (i.e. feed attempts), there was no significant interaction between species and turbidity (negative binomial GLM: species * (scale(turbidity)): LRT_1,66_ = 0.02, *P* = 0.88) or a linear effect of turbidity (negative binomial GLM: (scale(turbidity)): LRT_1,67_ = 1.18, *P* = 0.27). The effect of species was significant, *O. niloticus* attempted to feed on a greater number of food items (negative binomial GLM: species: LRT_1,67_ = 63.77, *P* < 0.001). Adding a quadratic term for turbidity, however, resulted in a significant interaction between turbidity and species (negative binomial GLM: species * poly(scale(turbidity)): LRT_2,64_ = 11.1, *P* = 0.003). This interaction and the trends in Fig. 3a suggest that the number of feed attempts by *O. amphimelas* was the greatest at the intermediate turbidity, but there was no such quadratic relationship in the trials with *O. niloticus*.

**Fig. 3.**
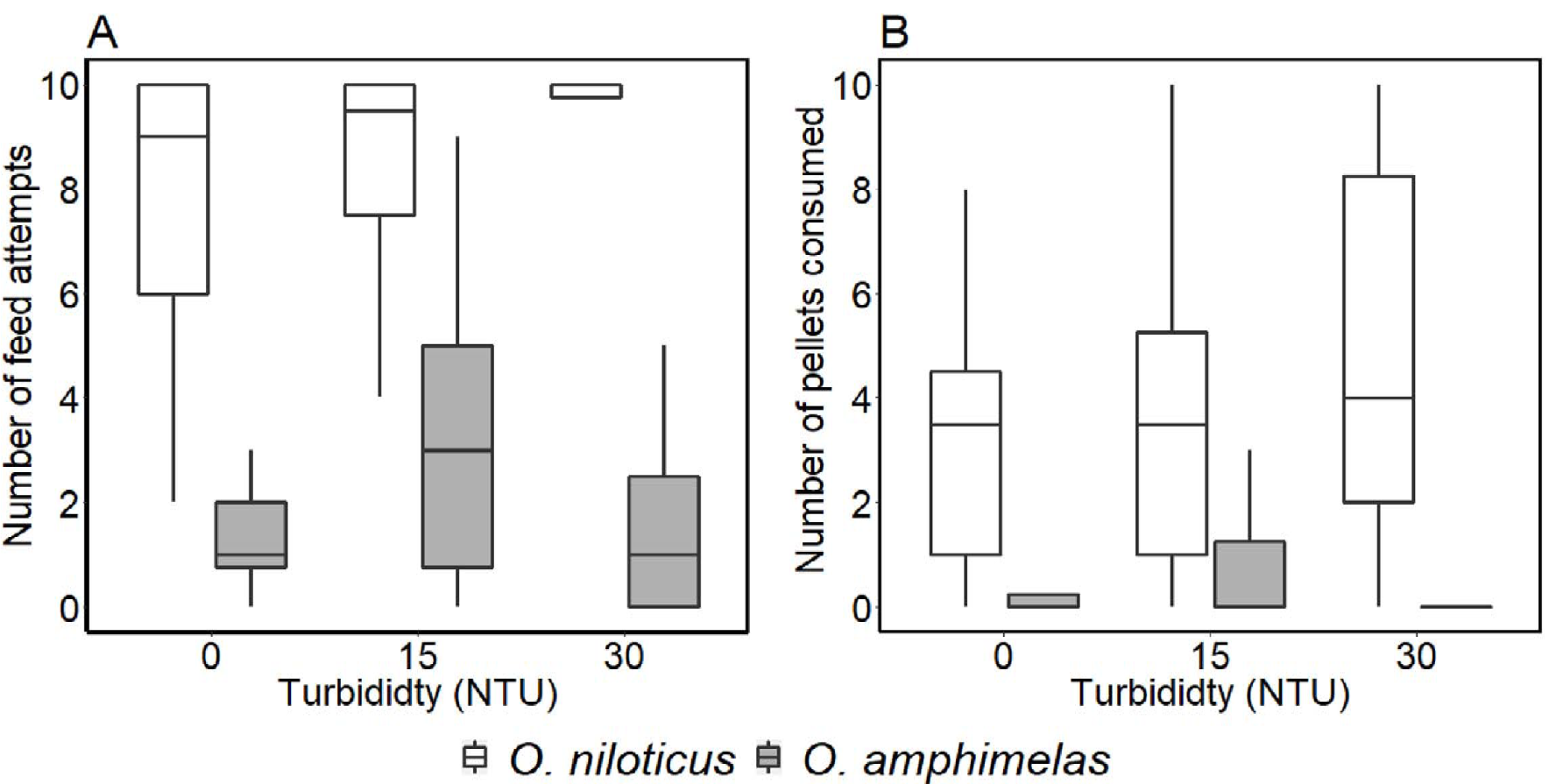
The number of (A) feed attempts and (B) pellets consumed across turbidity treatments for both species, *O. niloticus* (white) and *O. amphimelas* (grey). The boxes represent the interquartile range with the middle line displaying the median. Each whisker represents the position of 50% of values outside of the interquartile range.

No effect on total food consumption was found in the interaction of species and turbidity (negative binomial GLM: species * (scale(turbidity)): LRT_1,66_ = 1.19, *P* = 0.27; Fig. 3b). As main effects, turbidity did not affect how many pellets were consumed (negative binomial GLM: (scale(turbidity)): LRT_1,67_ = 3.29, *P* = 0.06) but *O. amphimelas* consumed less than *O. niloticus* (negative binomial GLM: species: LRT_1,66_ = 13.87, *P* < 0.001). Analysing the data by species separately, turbidity did not affect the number of pellets consumed by *O. amphimelas* (negative binomial GLM: (scale(turbidity)): LRT_1,32_ = 0.76, *P* = 0.38) or *O. niloticus* (negative binomial GLM: (scale(turbidity)): LRT_1,32_ = 1.36, *P* = 0.24).

### Experiment two

The likelihood of leaving the shelter for the first time did not differ between species (CPH: Hazard ratio (HZ) = 1.01, N = 244, *P* = 0.9). When analysing the data from the two species separately, the likelihood of leaving the shelter was not significantly different between turbidity treatments for either species (*O. amphimelas*: CPH: HZ = 0.68, N = 116, *P* = 0.06; *O. niloticus*: CPH: HZ = 0.73, N = 128, *P* = 0.1; Fig. 4 a, b). The likelihood of crossing the midline for the first time did not differ between species (CPH: HZ = 0.68, N = 244, *P* = 0.1). However, the likelihood of crossing the midline was significantly lower in high turbidity (15 NTU) than in clear water for both species (*O. amphimelas*: CPH: HZ = 0.51, N = 116, *P* = 0.001; *O. niloticus*: CPH: HZ = 0.57, N = 128, *P* = 0.003; Fig. 4 c, d).

**Fig. 4.**
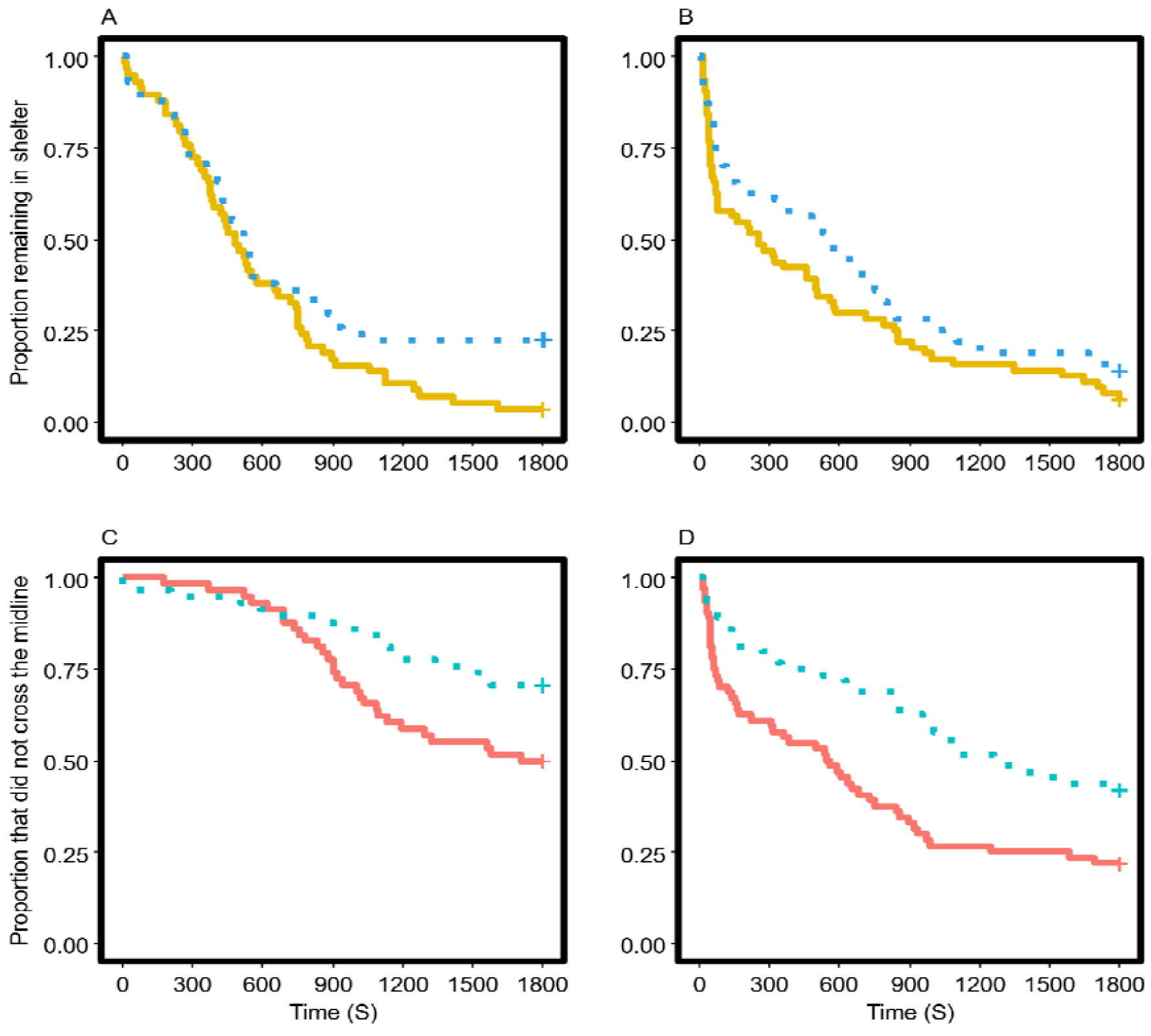
Kaplan-meier event estimates displaying the proportion of individuals to remain in the shelter (A) *O. amphimelas* and (B) *O. niloticus* and the proportion that did not cross the midline (C) *O. amphimelas* and (D) *O. niloticus* at 0 (solid) and 15 (dotted) NTU.

No effect on total time outside shelter was found in the interaction of species and turbidity (GLMM: species * (scale(turbidity)): □^2^_1_ = 0.33, *P* = 0.5). As main effects the total time outside of the shelter did not differ between species (GLMM: species □^2^_1_ = 0.53, *P* = 0.4; Fig. 5), and was lower in turbid water for both *O. amphimelas* (GLMM: scale(turbidity) □^2^_1_ = 10.85, *P* = 0.0009; Fig. 5) and *O. niloticus* (GLMM: turbidity □^2^_1_ = 10.23, *P* = 0.001; Fig. 5).

**Fig. 5.**
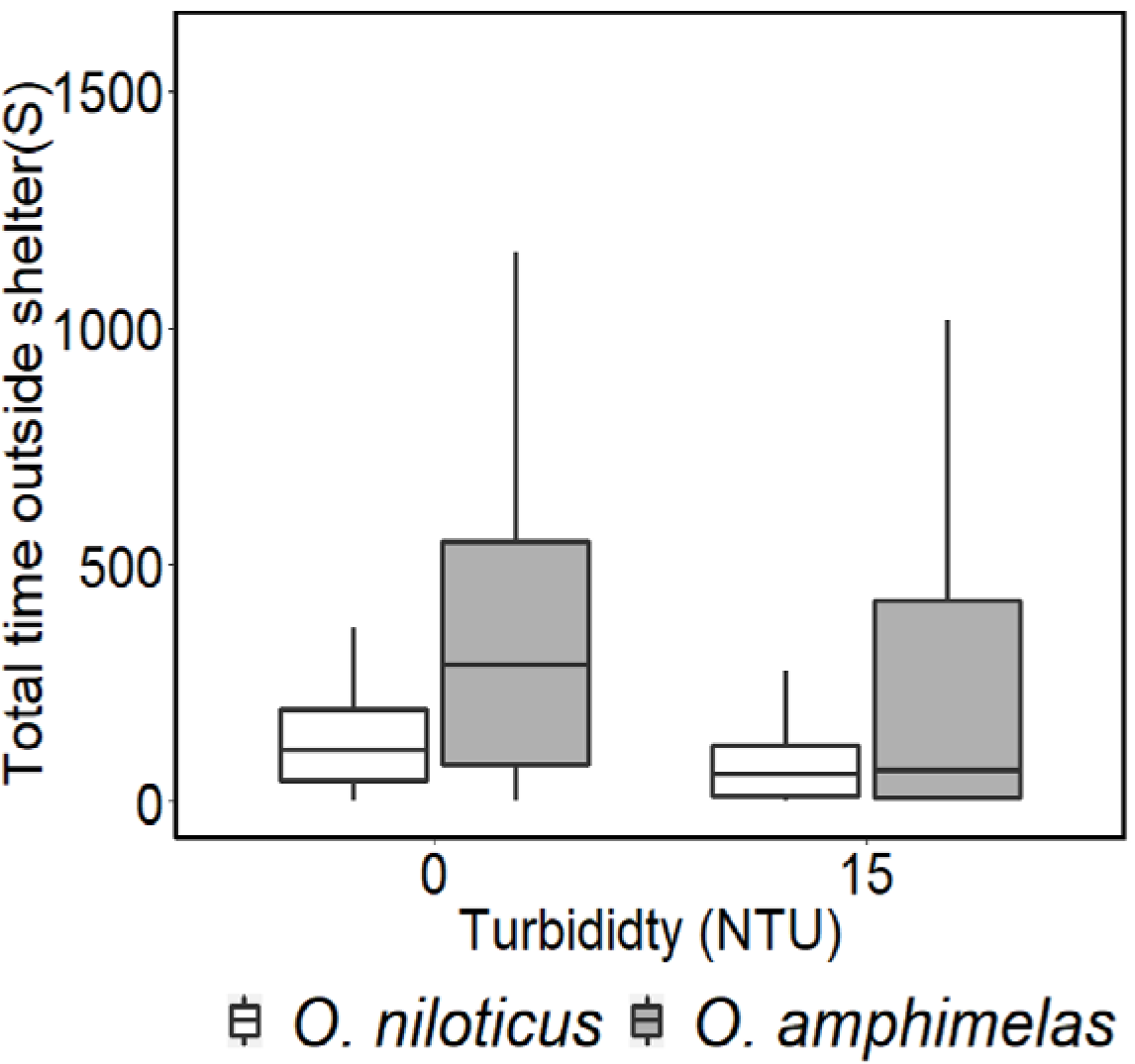
Total time spent outside the shelter. The median are horizontal lines within the boxes, the interquartile range is the area within the boxes and the whiskers display the data within 1.5 x IQR.

Spearman rank correlation coefficients showed inter-individual consistency in the latency of *O. amphimelas* to first leave the shelter in the 15 NTU treatment (N = 29, *r*_*s*_ = 0.405, *P* = 0.029) but all other correlation coefficients were nonsignificant (Table 1), suggesting little support for consistent inter-individual variation in these species.

**Table. 1.**
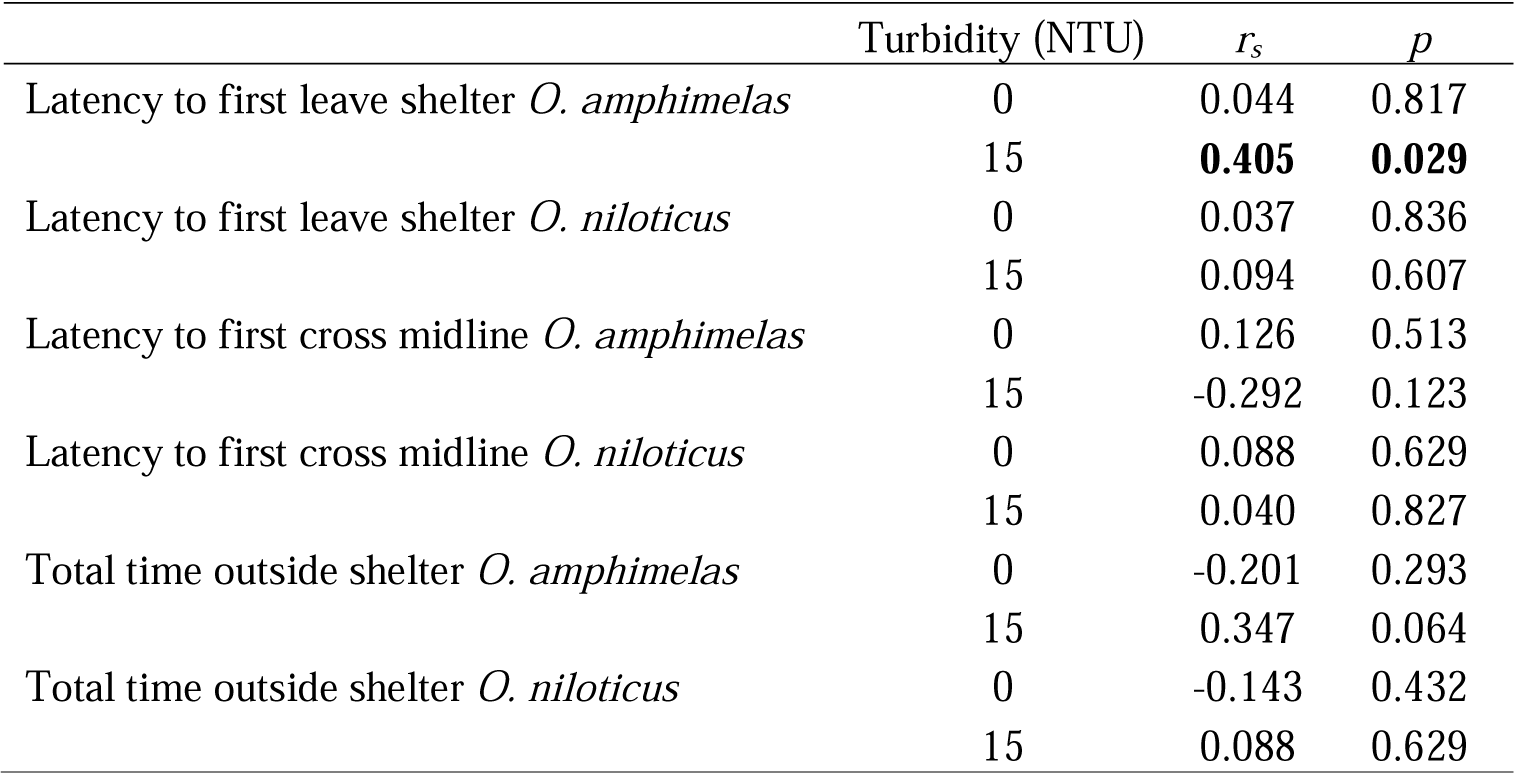
Spearman rank correlation coefficients (*r*_*s*_) for behaviours measured in two repeated trials in clear and turbid water. 29 *O. amphimelas* and 32 *O. niloticus* were tested. Significant values at 0.05 are shown in bold

## Discussion

Our results show that *O. niloticus* from aquaculture stock, which are those most likely to be introduced outside of the species’ native range, started to feed more quickly, encountered more food items and consumed more in total than *O. amphimelas* bred from wild-caught parents. The latency to first feed by *O. amphimelas* was unaffected by turbidity, while *O. niloticus* fed more readily when the water was turbid (15 and 30 NTU) than when it was clear (0 NTU). The number of feed attempts made by *O. niloticus* was not affected by turbidity level, while *O. amphimelas* had the highest number of feed attempts at the intermediate turbidity of 15 NTU. This peak at intermediate turbidity was not found in the number of pellets consumed, which did not depend on turbidity for either species. In experiment two, the time taken to first leave the refuge was unaffected by turbidity but fish showed reduced risk taking by being slower to cross the midline of the arena and spent less time overall outside of the shelter. This suggests that the difference between the species in their foraging response to turbidity in experiment one was not a result of differences in risk-taking tendency (boldness). There was little evidence of consistent differences between individuals (i.e. personality variation) in either clear or turbid water. *O. amphimelas* did display a positive correlation over different days in the latency to first leave the shelter in the turbid treatment, suggesting some indication of a consistent shy-bold continuum in turbid water. However, the lack of similar results across other behaviours makes this only suggestive in *O. amphimelas* and shows that personality variation in boldness is weak in these species compared to other fish (Brown et al. 2005; Biro et al. 2010; Bevan et al. 2018; Ehlman et al. 2019; Szopa-Comley et al. 2020).

Turbidity can have a negative impact on visual foragers by increasing foraging latency, reducing reaction distance and reducing attack success; however our study showed that the feeding latency of *O. niloticus* was actually lower in the turbid treatment (Gregory and Levings 1996; Turesson and Brönmark 2007; Becker et al. 2016). *O. niloticus* may compensate for reduced visual information in turbid environments through increased activity and/or increasing reliance on another sensory modality. Past studies on *O. niloticus* demonstrate it has an acute sense of olfaction that is used for foraging when vison is limited (Marusov and Kasumyan 2017). This could explain the ability of *O. niloticus* to forage in turbid water, as species dependent on olfaction are not affected when foraging in turbid environments (Lunt and Smee 2015). Compensatory factors such as these may explain why the time to first feed by *O. amphimelas* was not affected by turbidity, but does not explain the more rapid time to feed in turbid water by *O. niloticus*, or the improvement in the number of feeding attempts by *O. amphimelas* from the clear to the intermediate turbidity treatment. The perception of risk by *O. niloticus* may instead have been lower in turbid water, possibly because turbidity was used as a refuge (Lehtiniemi et al. 2005; Ferrari et al. 2014), although this was not supported by the results of experiment two, which showed reduced risk taking in both species in turbid water. Our results may indicate that the response to turbidity may be highly context specific, having a different effect depending on the presence or absence of food.

For *O. amphimelas*, intermediate levels of turbidity may be advantageous when foraging but do not increase the motivation to initially forage as there was no effect of turbidity on the latency to first feed. This may allow *O. amphimelas* to forage in moderately turbid environments and potentially mitigate the effects of anthropogenic turbidity. However, our results indicate this is only effective up to a certain threshold between 15 and 30 NTU. As previous work has indicated that turbidity between 26 and 141 NTU reduces the diversity and volume of stomach contents of fish species (Stuart-Smith et al. 2004), establishing at what range feeding efficiency is negatively affected is important and could help maintain populations of native species.

Considering the role *O. niloticus* plays in increasing turbidity, its presence may initiate a reduction in quality of clear water habitats or encourage a further decline in those already degraded (Zhang et al. 2017). Our study shows that some aspects of feeding by *O. niloticus* increases in turbid water suggesting that habitat degradation driven by *O. niloticus* can be beneficial to its foraging. This follows previous studies that suggest that degraded habitats are favourable to invasive generalists allowing colonisation where native species decline (Marvier et al. 2004). If an introduction of *O. niloticus* leads to increased turbidity and so improves its feeding efficiency, it would become even more adept at foraging than native species. Thus, the introduction of *O. niloticus* could facilitate a positive feedback loop that results in it dominating competition with native species (Mainka and Howard, 2010). This is especially relevant considering the foraging threshold of turbidity seen in *O. amphimelas*. If *O. niloticus* is introduced to an area where populations of *O. amphimelas* reside it may increase the level of turbidity above this threshold potentially leading to the decline in foraging efficiency of *O. amphimelas*.

Our results demonstrate how adept *O. niloticus* from aquaculture stock is at foraging even compared to a closely related species in the same genus, and that high levels of turbidity can constrain foraging in *O. amphimelas* by reducing the number of encounters with food, while increasing the perception of risk in both species. Therefore, the presence of *O. niloticus* presents a threat to *O. amphimelas* and very likely other native cichlid species where it has been introduced, as it has the potential to increase turbidity and then excel at foraging within this degraded environment. Preventing habitats from becoming turbid is a potential management tool in combating the spread and negative effects of *O. niloticus* and similar invasive species. However, despite turbidity influencing the foraging behaviour of both species, we found no effect of turbidity on total food consumption, and food consumption was much greater in *O. niloticus* at all levels of turbidity. Further observations of *O. niloticus in situ* are required to further understand how their behaviour can affect native species across its wide global distribution, with respect to foraging and other behaviours (Linde et al. 2008a, b; Salazar Torres et al. 2016; Zhang et al. 2017; Champneys et al. 2018; Champneys et al. 2020). This is especially important if other native species exhibit a lower threshold of turbidity tolerance than *O. niloticus*.

## Acknowledgements and Funding

We thank Professor George Turner from Bangor University for providing us with the *O. amphimelas* for this study. T.S.C. was funded by a NERC GW4+ FRESH CDT PhD studentship (NE/R011524/1). C.C.I. was supported by the Natural Environment Research Council, grant number NE/P012639/1.

## Declarations

### Ethical approval

All experimental procedures and housing conditions were in accordance with the ethical standards of the University of Bristol and ethical approval was granted by the University’s Animal Welfare and Ethical Review Body (UB/18/067).

### Consent to participate

Not applicable.

### Consent for publication

Not applicable.

### Availability of data and material

All data generated or analysed during this study are included in this published article (and its supplementary information files).

### Code availability

The code used in the current study is available from the corresponding author on reasonable request.

